# Universal baleen whale microsatellite panel for individual identification and power to detect parentage

**DOI:** 10.1101/2023.04.12.536337

**Authors:** Marcos Suárez-Menéndez, Martine Bérubé, Lutz Bachmann, Peter Best, Mads Peter Heide-Jørgensen, Veronique Lesage, Tom Oosting, Rui Prieto, Christian Ramp, Jooke Robbins, Richard Sears, Mónica A. Silva, Marc Tollis, Els Vermeulen, Gísli A. Víkingsson, Øystein Wiig, Per J. Palsbøll

## Abstract

Highly polymorphic single tandem repeat loci (STR, also known as microsatellite loci) remain a familiar, cost efficient class of markers for genetic analyses in ecology, behavior and conservation. We characterize a new universal set of ten STR loci (from 28 potential candidate loci) in seven baleen whale species, which are optimized for PCR amplification in two multiplex reactions along with a Y chromosome marker for sex determination. The optimized, universal set of STR loci provides an ideal starting point for new studies in baleen whales aimed at individual-based and population genetic studies, and facilitates data sharing among research groups. Data from the new STR loci were combined with genotypes from other published STR loci to assess the power to assign parentage (paternity) using exclusion in four species: fin whales, humpback whales, blue whales and bowhead whales. We argue that parentage studies should present a power analysis to demonstrate that the specific data are sufficiently informative to assign parentage with statistical rigor.

## Introduction

Conservation genetic assessments typically draw inferences from the degree of genetic diversity within and among populations or individuals. Estimating the degree of relatedness among individuals can yield insights into the social system, mating strategy and population structure of endangered species. During recent years, single nucleotide polymorphisms (SNPs) have gained in popularity relative to short tandem repeat loci (STR, also known as microsatellites (Tautz 1989), due to their greater abundance in genomes (Morin, Luikart, and Wayne 2004). However, genotyping STR loci in conservation is still common, and often the only viable alternative when resources are scarce or the research objective requires genotyping large sample sizes and highly polymorphic loci. The specific set of STR loci genotyped is often identified and characterized in the targeted species or from published resources in closely related species. As a result, different combinations of STR loci are genotyped among individual laboratories and studies (with some notable exceptions, such as salmonids, Moran et al. 2006). Although there are obvious downstream advantages to genotype “universal” sets of STR loci, such optimization is tedious, due to experimental issues (e.g., cross-species amplification, null alleles and ascertainment bias Primmer et al. 1996) and varying levels of polymorphism among species. However, the DNA sequences flanking STR loci are often conserved among closely related taxa, which facilitates genotyping of homologous loci in closely related species. We developed a set of ten STR loci and a sexing marker that can be genotyped in seven baleen whale species in two multiplex polymerase chain reactions (PCR, Saiki et al. 1986). Identifying and optimizing cross-species amplifying STR loci would not only facilitate the development of new studies, but also enable data sharing among laboratories and the re-use of published data.

A main use of STR loci is the identification of related individuals in a population (Blouin 2003). Employing different numbers of loci and the varying level of polymorphism among loci has a stark effect on the power to infer accurate relationships between individuals. For example, individuals that are closely related to a putative offspring (e.g. siblings or its own offspring) may incorrectly be assigned as a parent if the number of STR loci is insufficient given the sample size. Most parentage assessments in wild populations studies were aimed at terrestrial mammals, such as brown bears (*Ursus arctos*, Shimozuru et al. 2022) or animals in captivity, such as the cultured giant groupers (*Epinephelus lanceolatus*, Weng et al. 2021). In *Mysticeti* (baleen whales), a few parentage studies have been performed in humpback whales (*Megaptera novaeangliae*, Cerchio et al. 2005; Cypriano-Souza et al. 2010; Clapham and Palsbøll 1997; Nielsen et al. 2001), minke whales (*Balaenoptera acutorostrata*, Skaug, Bérubé, and Palsbøll 2010), North Atlantic right whales (*Eubalaena glacialis*, Frasier et al. 2007) and southern right whales (*E. australis*, Carroll et al. 2012). Most studies do not assess the informativeness of the STR loci employed prior to performing parentage analysis and thus have little or no insight into the error rate (i.e., false positives). Here we present a procedure (and supply the associated code) to determine how many STR loci are needed for a rigorous assignment of parentage using parentage exclusion probabilities. Towards this specific goal, we selected STR loci from published sources and characterized 28 new STR loci, which we employed with previously published STR loci to assess the number of STR loci required for rigorous parentage assignment in fin whales (*Balaenoptera physalus*), humpback whales, blue whales (*Balaenoptera musculus*) and bowhead whales (*Balaena mysticetus*).

## Materials and Methods

All tissue samples were collected as skin biopsy from free-ranging whales as described by Palsbøll et al. (1991). Samples were stored at -20 or -80 degrees Celsius (℃) in saturated NaCl with 25% dimethylsulphoxide (Amos and Hoelzel 1991).

### Molecular analyses

Total-cell DNA was extracted using either standard phenol-chloroform extractions (Sambrook and Russell 2001) or QIAGEN DNEasy^TM^ extraction columns for animal tissue, following the manufacturer’s instruction (QIAGEN Inc.). The quality of the DNA was assessed visually by electrophoresis through 0.7% agarose gel and the amount quantified with a Qubit^TM^ following the manufacturer’s instructions (Thermo Fisher Scientific Inc.). DNA extractions were adjusted to a final concentration at 10ng DNA/μL.

Candidate tetramer STR loci with the repeat motif GATA were identified using the software SCIROKO (ver. 3.4, Kofler et al. 2007) and the humpback whale genome assembly (Tollis et al. 2019). The search was conducted with the parameter settings: search mode (perfect repeats), minimum number of repeats (5), upper and lower bound of motif length (4), SSR-couple considerations (all).

Oligo-nucleotides for PCR amplification of each STR locus were designed using PRIMER3PLUS (ver. 3.2.6, Untergasser et al., 2012) with default parameter settings, except fixing the annealing temperature at 57 ℃ and the oligo-nucleotide length at 21 nucleotides (Table 1). For each pair of oligo-nucleotides, the forward oligo-nucleotide was extended with either a universal T7 or M13 DNA sequence (Schuelke 2000). The T7/M13 extension facilitated labeling of the amplification products with a fluorophore (6-FAM or HEX) of the complementary T7/M13 primer during PCR and hence detection during capillary electrophoresis on ABI 3730 Genetic Analyzer (Applied Biosystems Inc.).

**Table 1.**
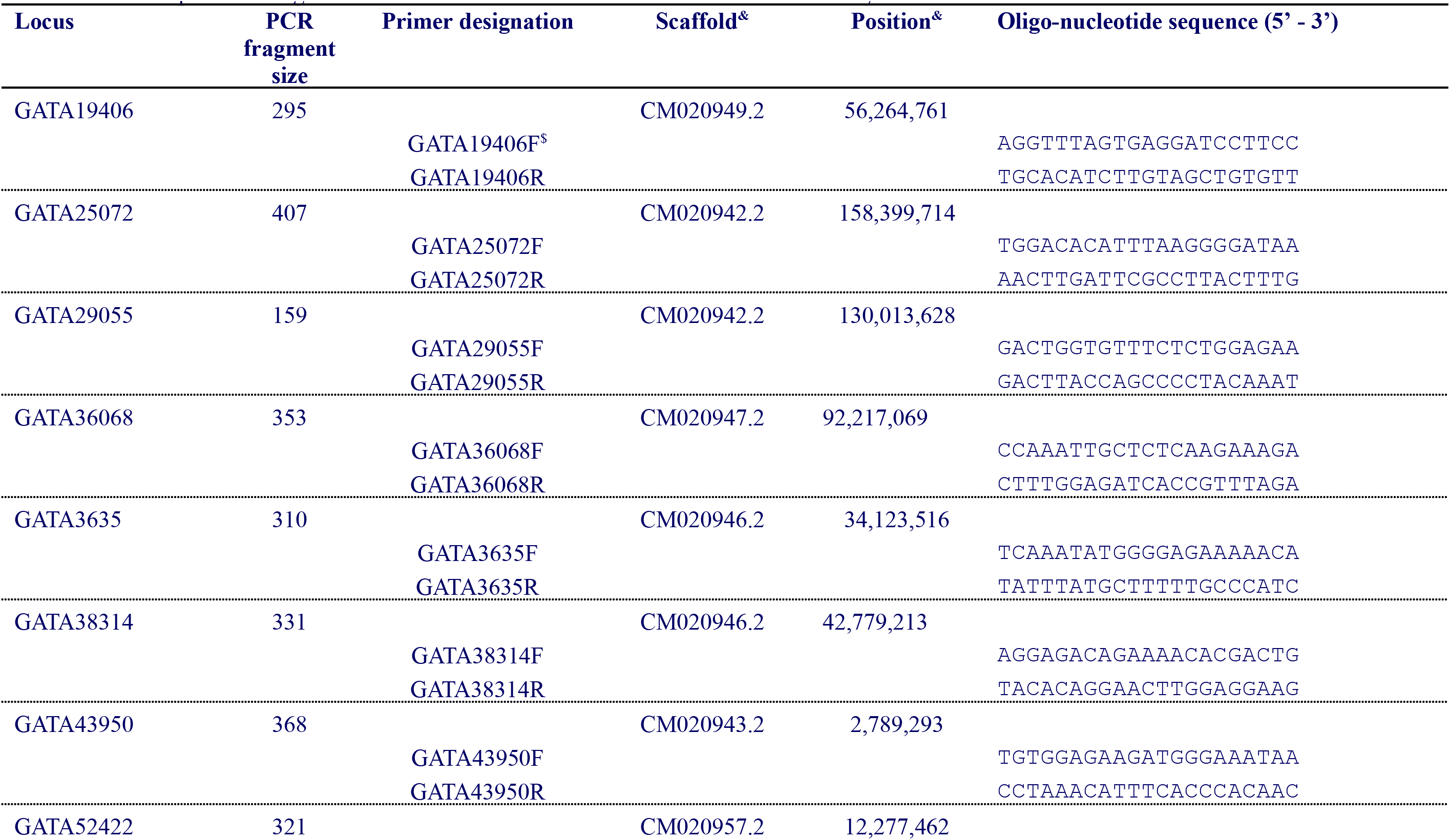
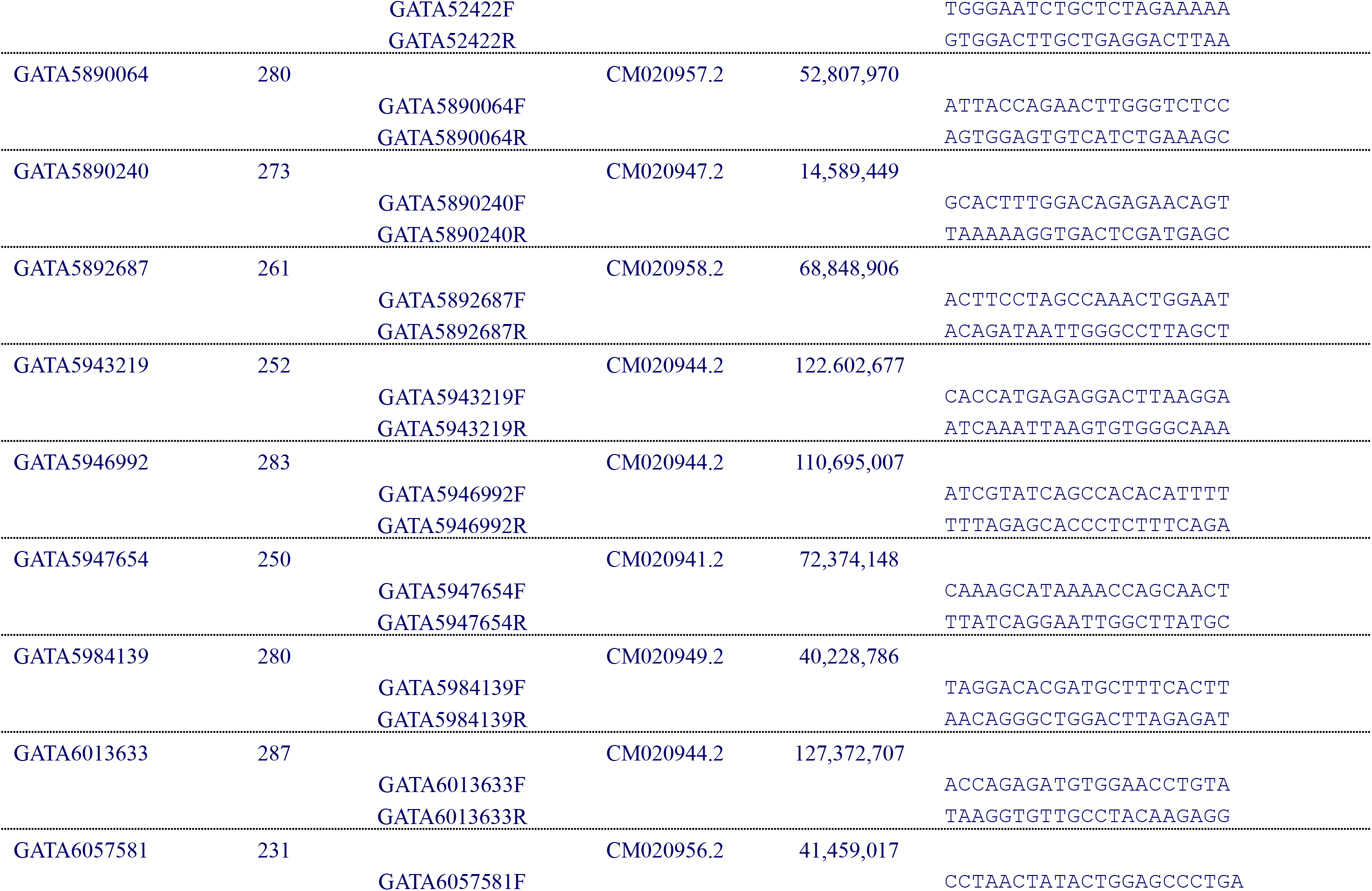
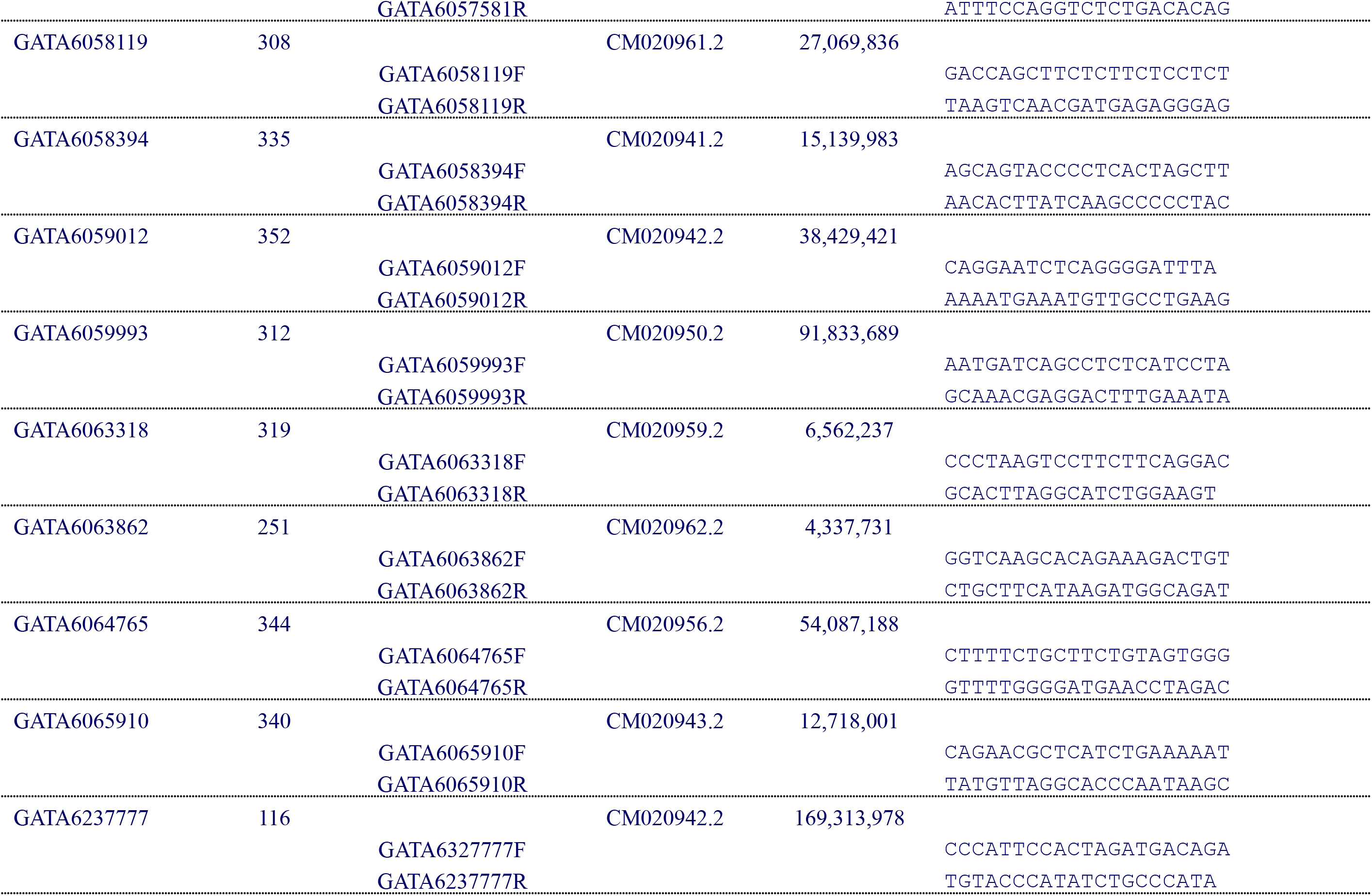
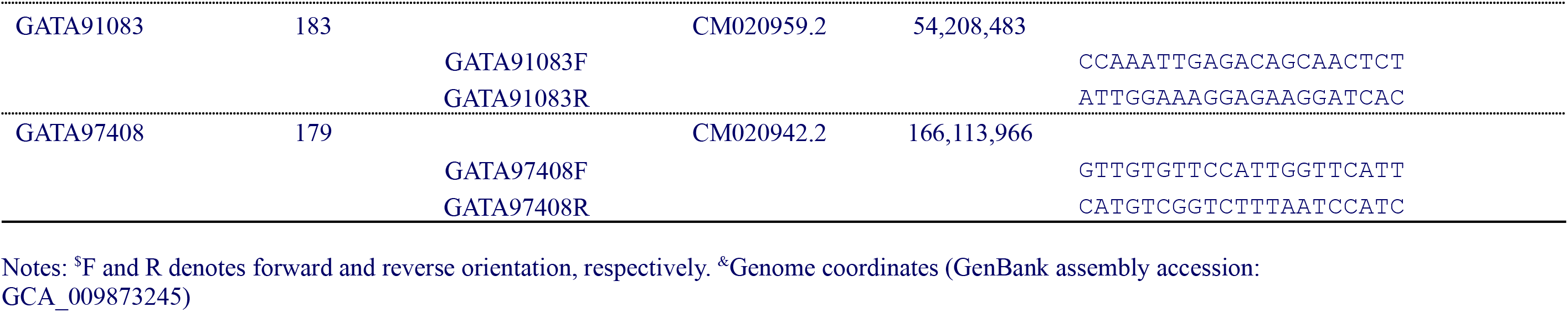
Primer sequences and genomic coordinates of the 28 new STR loci used in the study.

The genomic coordinates of each STR locus were determined by aligning the oligo-nucleotides against the reference blue whale genome at the chromosome-level scaffolds (Bukhman et al. 2022) using BOWTIE2 (ver 2.3.5.1, Langmead and Salzberg 2012) with the parameter settings defined by the preset *–very-sensitive*. PCR amplification were conducted under the following conditions in 10μL volumes, each with 10ng of genomic DNA, 67mM Tris-HCl, pH 8.8, 2mM MgCl_2_, 16.6mM (NH_4_)_2_SO_4_, 10mM *β*-mercaptoethanol, 0.2mM dNTPs, 1µM of each oligo-nucleotides as well as 0.4 units of *Taq* DNA polymerase (New England Biolabs Inc.). PCR reactions were subjected to two minutes at 94 °C followed by 35 cycles each with 30 seconds at 94 °C, 90 seconds at 57 °C and 30 seconds at 72 °C; followed by 10 minutes at 68 °C. The initial quality of the PCR amplification products was first assessed by gel electrophoresis in 2% agarose and 1xTBE at 175 volts for 25 minutes. After staining with ethidium bromide, amplification products were visualized under UV light at 260 nm. All candidate STR loci were amplified by as described above except for the three oligo-nucleotides which were added in the following concentrations: 1µM of the unlabeled locus-specific oligo-nucleotides, 0.5μM of the 5’end-labeled (HEX or 6-FAM) oligo-nucleotide and 0.5μM of the M13/T7-extended (see Table S2), unlabeled locus-specific oligo-nucleotide. PCR reactions were subjected to two minutes at 94 °C followed by 10 cycles each consisting of 30 seconds at 94 °C, one minute at 57 °C, and 30 seconds at 72 °C followed by an additional 27 cycles each with 30 seconds at 94 °C, one minute at 55 °C and 30 seconds at 72 °C, followed by 10 minutes at 72 °C. The amplification products were separated by capillary electrophoresis on an ABI 3730 Genetic Analyzer (Applied Biosystems Inc.). The size of the amplification products was estimated using the size standard GeneScan™ 500 ROX™ dye size standard (Applied Biosystems Inc.) with GENEMAPPER ver 4.0 (Applied Biosystems Inc.). One of each pair of oligo-nucleotide pairs was labeled with either 6-FAM, HEX or NED for those candidate STR loci selected for further analysis (Table S1) and genotyped in 48 individuals from each of seven mysticete species; humpback whale, fin whale, minke whale, sei whale (*B. borealis*), blue whale, southern right whale and bowhead whale.

Multiplex PCRs were performed using the Qiagen™ Multiplex Kit Plus (Qiagen Inc.) in 5µL reaction volumes following the manufacturer’s recommendations. PCR reactions were subjected to two minutes at 94 °C, followed by 35 cycles each with 30 seconds at 94 °C, 90 seconds at 57 °C and 30 seconds at 72 °C, followed by 10 minutes at 68 °C. PCR amplification products were separated and detected by capillary gel electrophoresis on an ABI 3730 Genetic Analyzer (Applied Biosystems Inc.) including a size standard GeneScan^TM^-500 ROX (Applied Biosystems Inc.). The length of each PCR product was determined with GENEMAPPER™ (ver. 4.1, Applied Biosystems Inc.).

Finally, a random set of 60 DNA extractions collected from different individual fin, humpback, bowhead and blue whales were genotyped at 30, 32, 22 and 20 loci, respectively, and the sex determined by co-amplification of a Y chromosome specific marker (Table 2) for the parent and offspring (PO) and sire-dam-offspring trio (SDO) assignments.

**Table 2.** Multiplex oligo-nucleotide panels for mysticete species.

| **Panel A** |  |  |  |  |  |
| --- | --- | --- | --- | --- | --- |
| Locus | Primer<br>Reverse | Primer<br>Forward | Dye | Range |  |
| GATA6237777 | R | F | HEX | 108 | 152 |
| GATA6064765* | R2 | F2 | HEX | 182 | 230 |
| GATA6063318 | R | F | HEX | 300 | 350 |
| GATA91083 | R | F | FAM | 149 | 220 |
| GATA38314 | R | F | FAM | 303 | 367 |
| **Panel B** |  |  |  |  |  |
| Locus | Primer<br>Reverse | Primer<br>Forward | Dye | Range |  |
| GATA5947654* | R3 | F3 | HEX | 130 | 160 |
| GATA5984139 | R | F | HEX | 204 | 280 |
| GATA52422RF | R | F | HEX | 320 | 479 |
| GATA25072* | R2 | F2 | FAM | 76 | 148 |
| GATA97408 | R2 | F2 | FAM | 164 | 215 |
| SRY | R | F | FAM | 332 | 332 |

\*New primers were re-designed in order to get different size fragments to fit the multiplex: GATA25072F2 (5'-CACCTGCTTTAAACTGTGTATAGT-3'), GATA25072R2 (5'-GATCTAGCAACTCTTTCTAGGC-3'), GATA6064765F2 (5'-GTACAAATGCACTTTCTCCCG-3') and GATA6064765R2 (5'-AGGCACTTATCAGTTCCAAGT-3'), SRYF, (5'-TGTGAACGGTGAGGATTA-3') and SRYR, (5'-GTGCATGGCTCGTAGTCT-3') GATA5947654R3 (5'-TCAGCCTCCATAATTGCATAAG-3') and GATA5947654F3 (5'-GTTATTAGATAGGGTTCTCTGCAG-3'), GATA97408F2 (5'-CTCCCACCACTGATTTGTAATA-3') and GATA97408R2 (5'-AGCCTAGTTTTATGGTACCTCT-3').

### Data analysis

Allele frequencies, the observed (*H*_O_) and expected (*H*_E_) heterozygosity, as well as the probability of identity for pairs of unrelated individuals (P(*I*), Paetkau and Strobeck 1994) corrected for low sample sizes were implemented in GIMLET (ver. 1.3.3, Valière 2002). The deviations from the expected Hardy-Weinberg genotype frequencies (HW) was estimated using GENALEX (ver. 6.5, Peakall and Smouse 2012; 2006). The non-exclusion probability of the first parent (*P*_NON-EXCL_, i.e., the probability of failing to exclude an unrelated, non-parent, Selvin 1980) was estimated using CERVUS (ver. 3.0.7, Kalinowski, Taper, and Marshall 2007).

### Parent-offspring assignment

PO pairs were identified using custom Python (ver. 3) scripts (see Data availability). The scripts identify all pairs of multilocus genotypes that segregate according to Mendelian expectations for parent and offspring. SDO trios were identified in a similar manner, i.e., by first identifying putative dam and offspring pairs among all possible pairs of multilocus genotypes requiring at least one sample to be from a female. Subsequently, putative sires were identified among male genotypes that were consistent with the putative dam and offspring genotypes and Mendelian segregation. The assessment did not allow for genotyping errors, i.e., putative parent and offspring pairs with loci that did not segregate according to Mendelian expectations were rejected.

KININFOR (ver. 2, Wang 2006) was employed to assess the informativeness of the different relatedness and relationship estimators. Since most relatedness estimators correlate (Table S4), the “informativeness of relationship” (IR) criterion was used to rank the STR loci in terms of their statistical power to discern among different degrees of relatedness. Two assessments were conducted to assess the power of a given number of STR loci to infer PO pairs. The first assessment was based upon the most or least informative STR loci (ranked by their IR value, see above); the second on a random sample of loci sampled among all loci genotyped and without replacement. In both assessments, data sets from eight to the maximum number of loci genotyped were generated and for each set of loci PO pairs and SDO trios identified as described above. The assessment of the effect of locus IR values on the ability to identify PO pairs and SDO trios was conducted in two ways, starting with the loci with the highest and lowest IR values, respectively.

### Assessments of paternity assignments

Close relatives to the putative offspring are most likely to be incorrectly assigned as the sire. Behavioral observations (e.g., Frasier et al. 2007) and genetic studies (e.g., Clapham and Palsbøll 1997) suggest that baleen whales are promiscuous and mating randomly. Although there is a possibility that a dam might mate with the same sire more than once and thus produce offspring that are related as full-siblings, such instances are likely rare unless the overall population size is very small. This implies that first (the offspring’s own offspring) and second order relatives (the offspring’s grandparents, grandchildren and half-siblings) are those individuals that are most likely to be incorrectly assigned as the sire. Since each offspring has more second order than first order relatives, we focused our assessment of the probability of incorrect paternity assignments on second order relatives. Using half-siblings as a representative of a second order relative, we assessed the probability of incorrectly assigning a half-sibling as the sire by generating simulated data sets during which *in silico* pairs of half-siblings were generated by randomly sampling alleles at each STR and sex locus from *in silico* sires and dams. The probability of an incorrect paternity assignment in each assessment was estimated as the proportion of half-siblings without any loci violating the Mendelian expectations of being a putative sire (allowing for a maximum of two Mendelian violations). The assessment was conducted with a maximum of 50 STR sampled at random, with replacement, from the observed loci. For each set of STR loci, 5,000 calves and half-siblings were simulated, and the assessment repeated 50 times. The median probability of an incorrect paternity assignment for each set of STR loci was smoothed using a Savitzky-Golay filter (Savitzky and Golay 1964) from which the “knee of the curve”, i.e. the point where adding additional STR loci had a diminished effect, was estimated using the Python package *kneed* (ver. 0.8.2, Satopaa et al. 2011).

## Results

### Newly characterized str loci

A total of 5,047 STR loci with the repeat motif GATA and minimum of five repeats was detected. We selected 28 STR loci which were deemed (a) of sufficient quality on the basis of the gel electrophoresis (above) and (b) likely polymorphic in mysticetes. These 28 selected STR loci were genotyped in eight DNA extractions in each species (Table 1). The majority of the STR loci amplified in the *Balaenopteridae* species, but less so within *Eubalaena* and *Balaena* (Table S1). Among the 28 candidate loci, ten STR loci that were polymorphic in most baleen whales were selected for further characterization. The ten candidate STR loci and a sex specific Y chromosome-specific locus were optimized for multiplexed PCR amplification. Amplification was conducted in two reactions; one with five STR loci and a second with five STR loci and the Y chromosome-specific locus (Table 2). In total, 48 DNA extractions from each mysticete species were genotyped with these two panels. Among the ten new loci, three loci were monomorphic in some species: GATA25072 (bowhead whale), GATA5947654 (southern right whale) and GATA6237777 (blue whale). *H*_E_ was similar among species (Table 3). The allele sizes at the loci GATA5947654 differed in the blue whale, and GATA5984139 differed in the bowhead whale and the southern right whale compared to the other species (Table S1). For the seven sets of 48 samples, the P(*I*) was estimated and ranged from 4.17x10^-3^ to 8.10x10^-1^. Samples with identical genotypes were inferred as duplicate samples. One pair of duplicates was found in the blue, sei, humpback and right whales. Only one sample was kept for the data analysis. The above aspects are tabulated in Table 3.

**Table 3.**
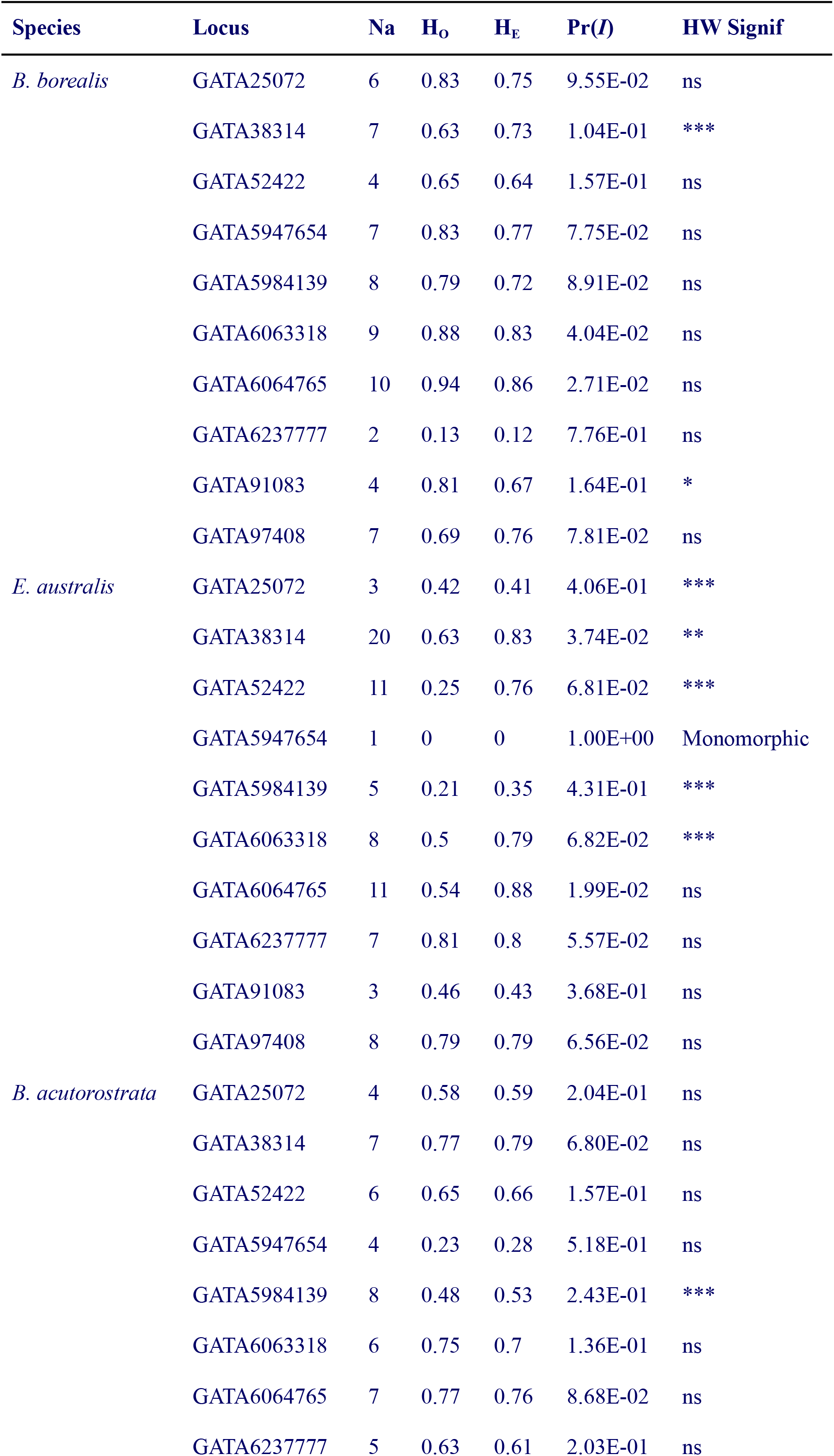
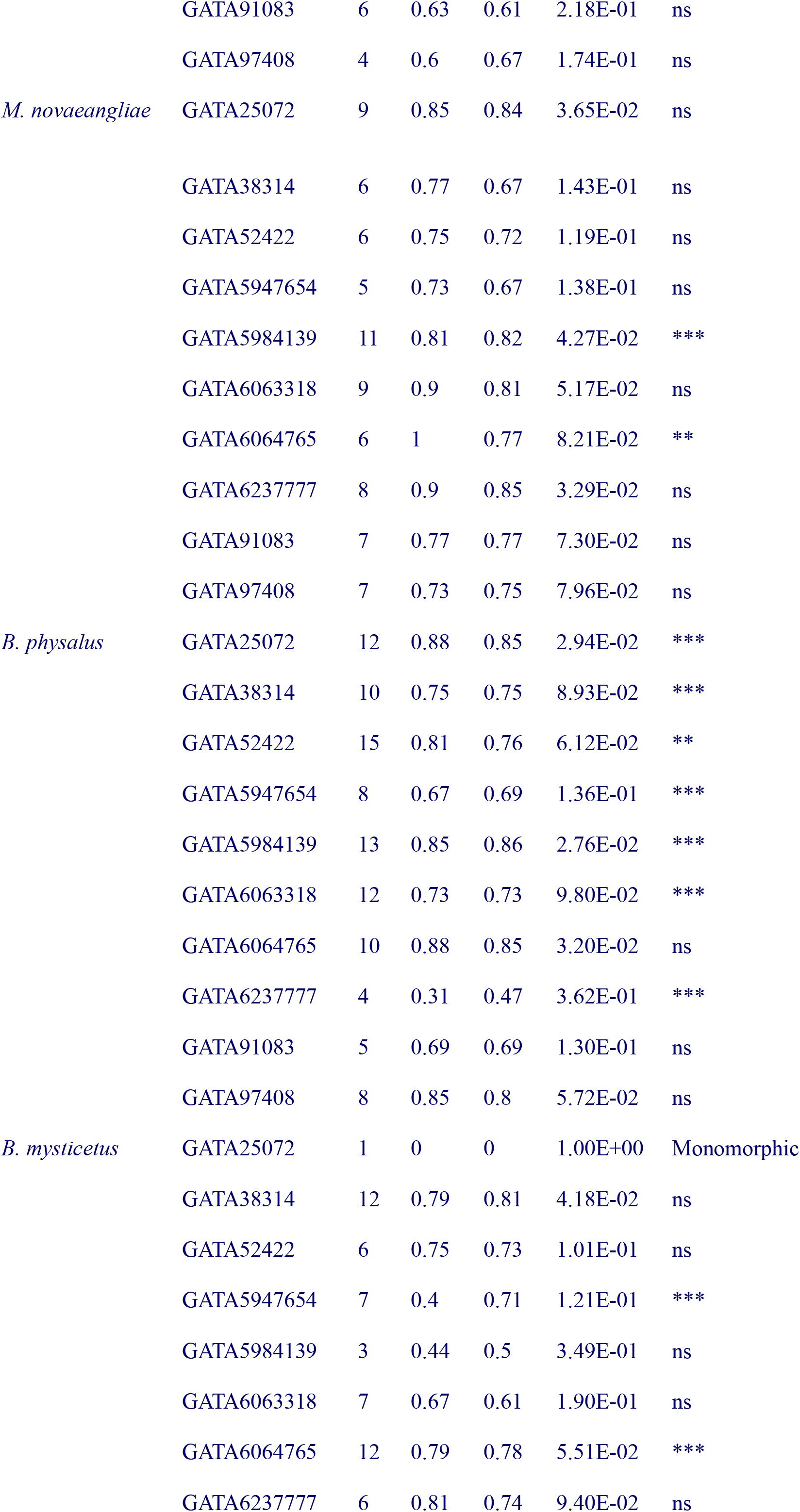
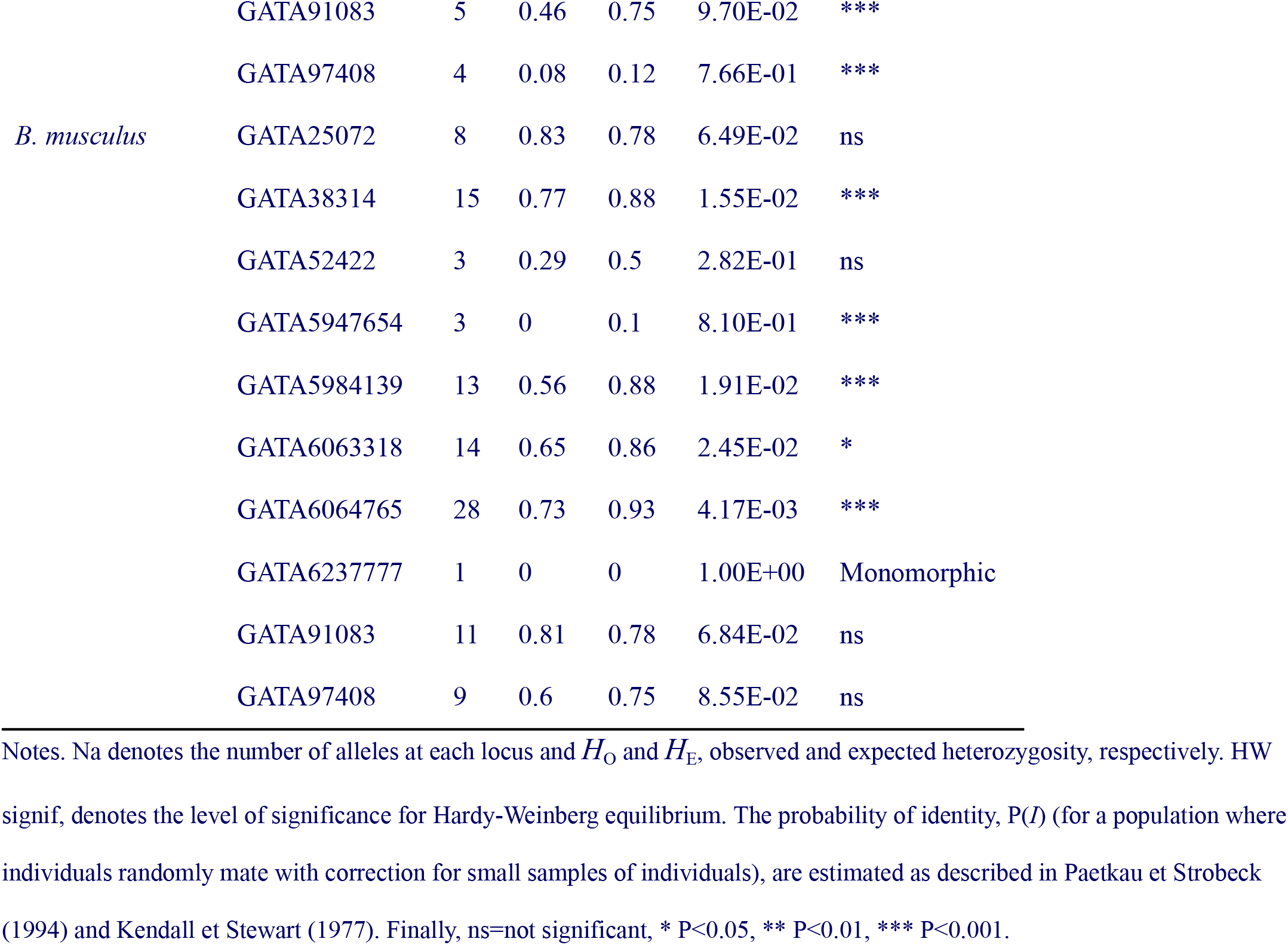
Summary statistics for 10 microsatellite loci in 48 individuals in a selection of mysticete species.

### Parentage assignments

In order to evaluate the impact of both the quantity and quality of genetic markers on the parentage inference, a total of 1,300 datasets were generated by selecting various combinations of loci from a dataset of 60 humpback whales individuals (30 females and 30 males) typed at 32 loci with no missing data. The selected 60 samples contained no duplicates.

*P*_NON-EXCL_ was estimated from samples of 60 randomly selected individuals in each of four species; fin, humpback, bowhead and blue whale (Table S2). The total of 32 STR loci genotyped in humpback whales represented a comparatively high level of polymorphisms with a mean locus *H*_E_ and *P*_NON-EXCL_ at 0.75 and 2.6x10^-7^, respectively. The same statistics were estimated for 30 loci in fin whales (mean locus *H*_E_ and *P*_NON-EXCL_ at 0.77 and 7x10^-8^, respectively), 20 loci in bowhead whales (mean locus *H*_E_ and *P*_NON-EXCL_ at 0.74 and 1.3x10^-5^, respectively) and 22 loci in blue whales (mean locus *H*_E_ and *P*_NON-EXCL_ at 0.68 and 6.2x10^-4^, respectively, Table S2).

PO pairs were identified among the 60 humpback whales genotyped at 32 STR loci as pairs of samples with the expected Mendelian segregation of alleles. If no missing data or mismatches were allowed, 17 PO pairs in total were detected. No additional (incorrect) PO assignments were detected if the assessment was based on 14 or more STR loci with the highest IR values, whereas a total of 27 loci among the least informative loci was required to achieve the same result (Table 4 and S3).

**Table 4.** Humpback whale combined exclusion probabilities of first parents estimated for ranked STR loci based on the informativeness of relationship criterion (IR).

| **Loci** | **Highest** | **Lowest** |
| --- | --- | --- |
| 8 | 0.002638 | 0.15063484 |
| 9 | 0.00151085 | 0.10668238 |
| 10 | 0.00087211 | 0.07395704 |
| 11 | 0.00050162 | 0.05130777 |
| 12 | 0.00028957 | 0.03557103 |
| 13 | 0.00017144 | 0.02442526 |
| 14 | 0.0001043 | 0.01612181 |
| 15 | 0.00006355 | 0.01068883 |
| 16 | 0.00003802 | 0.00676013 |
| 17 | 0.00002404 | 0.00404405 |
| 18 | 0.00001594 | 0.00246402 |
| 19 | 0.00001052 | 0.00149905 |
| 20 | 0.00000722 | 0.00088752 |
| 21 | 0.00000501 | 0.00051233 |
| 22 | 0.00000347 | 0.00029468 |
| 23 | 0.00000241 | 0.0001701 |
| 24 | 0.00000171 | 0.00009742 |
| 25 | 0.00000128 | 0.00005367 |
| 26 | 0.00000094 | 0.00002792 |
| 27 | 0.0000007 | 0.000014 |
| 28 | 0.00000053 | 0.00000685 |
| 29 | 0.00000042 | 0.00000338 |
| 30 | 0.00000034 | 0.00000154 |
| 31 | 0.00000029 | 0.00000065 |
| 32 | 0.00000026 | 0.00000026 |

Notes. Starting with the eight highest/lowest ranked loci.

The number of detected putative PO pairs using the minimum number of loci (eight) selected at random among the 32 loci varied between 34 to 104 (median 57). Both the median and range of the number of putative PO pairs declined with increasing number of loci. At 14 loci, a single of the 50 datasets yielded the same 17 PO pairs identified with the full data set (i.e., 32 loci). In this instance, the 14 loci at random happened to include all the most informative loci (as described above). At 24 or more loci, the median number of detected PO pairs was 17 PO (Figure 1, top panel).

**Figure 1.**
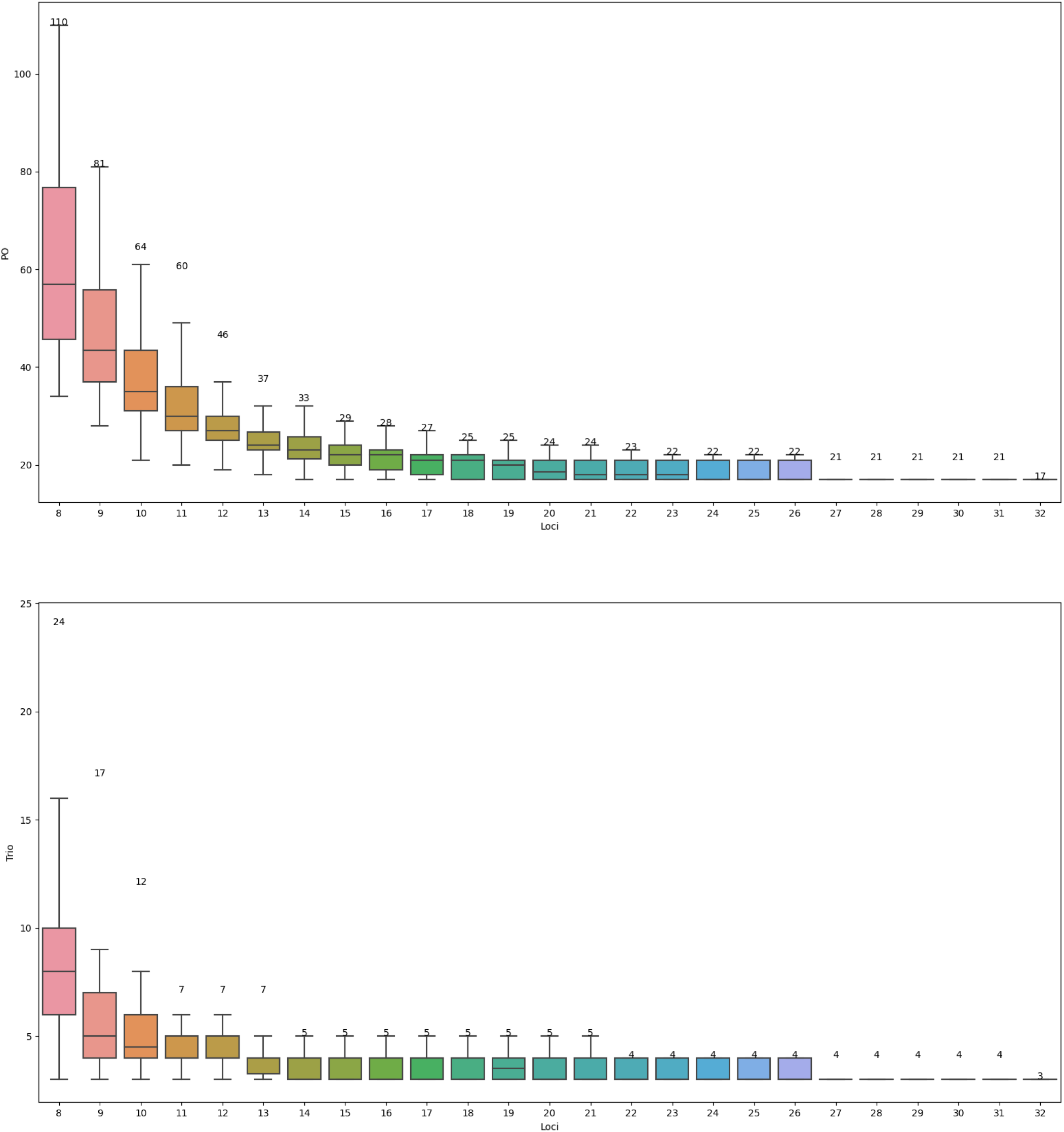
Effects of the number and informativeness (IR) of STR loci on parentage offspring and trio assignment. Notes. Values on top of each box indicate the maximum number of PO assignments in any dataset for that specific set of loci.

A similar pattern was observed when detecting SDO trios. Three SDO trios were identified among the 60 humpback whales in the full data set (32 STR loci). Datasets with only eight loci detected between three and 16 trios (median of eight) with an outlier at 24 trios. As expected the minimum of three trios was detected in all datasets. At 20 loci or higher, the median number of SDO trios detected was three (Figure 1, bottom panel).

The relative percentage of half-siblings assigned as sire was consistent with the combined exclusion probabilities for the different datasets in each species, with higher probabilities of incorrect assignments in blue whales and humpbacks compared to bowhead whales (Figure 2). The “knee” of the curve was at 15 and 17 loci at zero or one Mendelian violation in the bowhead whale; 15 and 17 in the fin whale; 16 and 17 in the humpback; and 15 and 17 in the blue whale. Allowing two Mendelian violations (or missing data) increased the critical number of loci to 21 in all but the fin whales for which it was 19. Allowing a few Mendelian violations (or missing data) had a significant impact on the probability of incorrect assignments, e.g., a 32 to 79-fold difference between zero and two Mendelian violations (or missing data) at 18 loci. The probability of an incorrect assignment approached zero at 24 loci in the fin whale, 26 in the humpback whale, 28 loci in the bowhead and at 30 loci in the blue whale.

**Figure 2.**
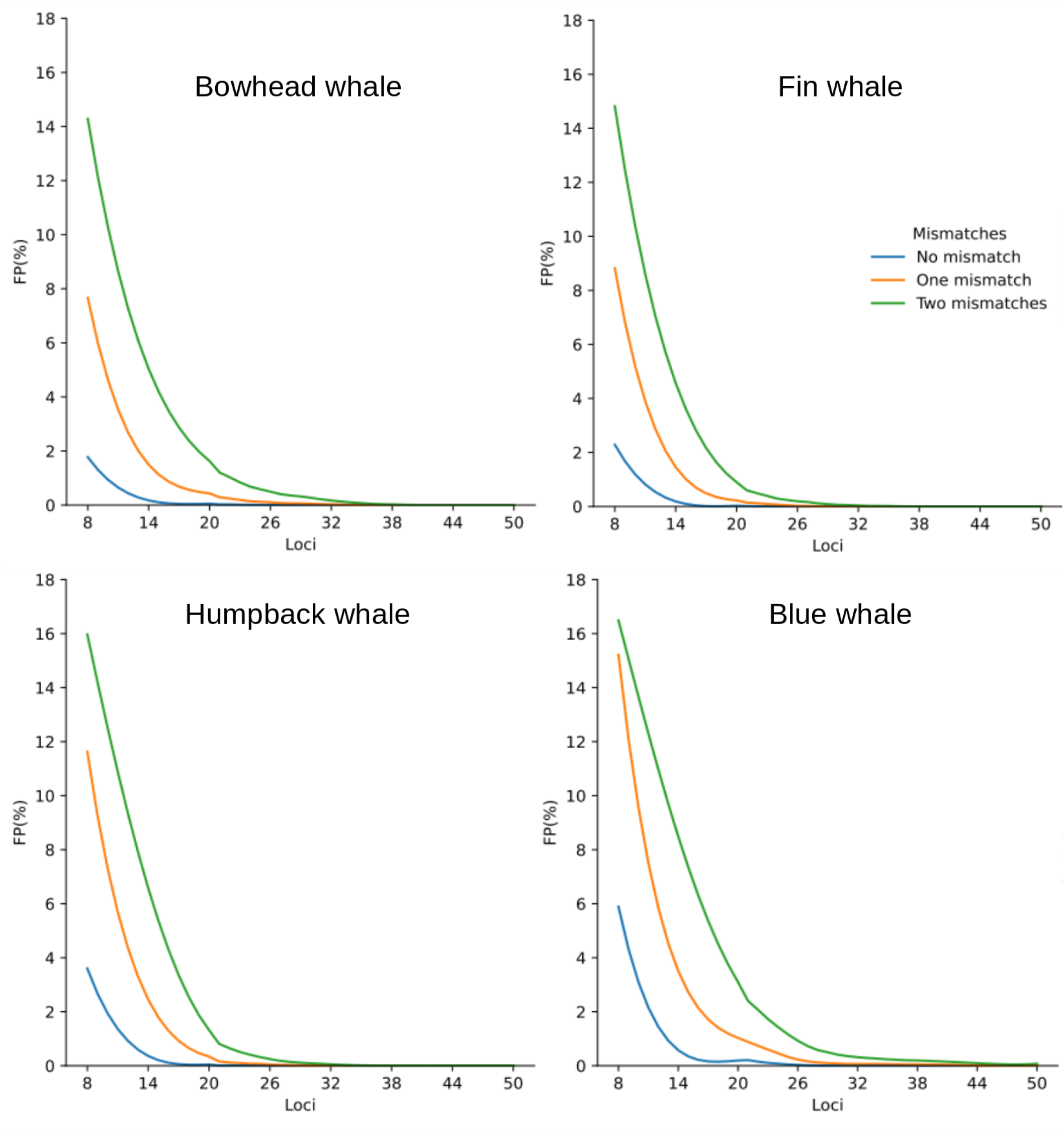
Percentage of false positives of paternity assignments per loci within each species.

## Discussion and conclusions

Multi-allelic STR loci remains a cost-efficient alternative to genomic approaches compared to whole genome sequencing and reduced representation methods, such as genotype-by-sequencing (e.g., Peterson et al. 2012). The low cost per sample makes STR genotyping an efficient means by which to screen large sample sizes for duplicate samples and pairs of first order relatives. PO pairs, and especially full pedigrees (i.e., SDO trios), are interesting for many purposes, such as kinship mark-recapture (Palsbøll 1999; Skaug 2001).

The increasing access to reference genomes further facilitates identification of STR loci in a cost effective manner. In this study, 28 “novel” STR loci were identified among ∼5,000 candidate STR loci in the humpback reference genome from which a genotyping assay comprising ten STR loci and a Y chromosome-specific locus were amplified in two multiplex PCR amplifications in most baleen whale species. Standardization of a basic set of STR loci applicable to a group of closely related species, simplifies initial genotyping within single research groups (incl. species identification as shown here) and enables sharing of datasets among laboratories working on the same species. Although the idea of developing multiplex panels to facilitate data sharing among laboratories working within the same group of species is not novel (Chambers et al. 2004) and has obvious long-term advantages, they remain rare (Beugin et al. 2017; Moran et al. 2006).

The second goal in this study was to facilitate the identification of PO pairs and SDO trios with a high level of confidence. The power of STR loci to discern between PO pairs and other close relationship categories is a function of the level of polymorphism at each included locus, the mating system of the targeted species, as well as reproductive rates. Some studies have inferred parentage employing only three STR loci (Zane et al. 1999) and among baleen whales assessments have been conducted with as few as nine to 13 STR loci (Cypriano-Souza et al. 2010; Carroll et al. 2012). In this study we demonstrate that the number of STR loci required to identify PO pairs with statistical rigor (using parentage exclusion probabilities) can be assessed in a relatively manner, hence maximizing the power of downstream parentage analysis. Unsurprisingly, we show that it is necessary to not only consider the number of STR loci but also the degree of polymorphism at each locus. The difference, in terms of loci to be genotyped to make rigorous inferences, were substantial between the loci with the lowest and highest IR values. Homologous STR loci differ in informativeness among species and populations, e.g., due to a high degree of relatedness among sampled individuals, which has a detrimental effect on the power to identify PO pairs and SDO trios. Thus, studies should be accompanied by a power analysis of the kinds shown here, for each specific dataset and reported when publishing the results. Currently only few parentage assessments undertake such a power analysis or accept high false positive rates (e.g., the 80% or 95% ”confidence” options available in CERVUS). Some studies reject PO pairs and SDO trios with two or more putative parent/sire, but accept PO pairs and SDO trios where only a single putative parent/sire was identified (e.g., Carroll et al. 2012; Frasier et al. 2007), thus ignoring the unsampled part of the population and the possibility that the identified parent/sire could be a non-parental close relative.

Genotyping errors might result in incorrect parentage/paternity exclusions and thus it is tempting to allow for a few Mendelian violations when assigning parentage/paternity. However, allowing a few Mendelian violations may result in a significant number of incorrect parentage/paternity assignments, especially when the number of loci or degree of polymorphism is low as shown here. As it is straightforward to assess the consequences of allowing some Mendelian violations on the rate of incorrect parentage/paternity assignments, the effect should be assessed and reported when interpreting the results. In the same vein, missing genotypes are common and often not reported or under-reported in many studies (e.g., Gerber et al. 2022). Putative genotyping errors are readily re-genotyped and checked. Consequently, the optimal approach is to check as many Mendelian violations as possible in the base data, rather than including such likely errors in the subsequent data analysis. Null-alleles (Paetkau and Strobeck 1995) could be the cause of loci violating Mendelian segregation, but are easily detected (both individuals homozygous for different alleles) and fixed (redesign PCR primers, and re-genotype all homozygous individuals).

Currently, missing genotypes are usually simply reported as the overall percentage of missing genotypes in the dataset, or as individuals with more than a minimum number loci. Since the effect of employing a reduced number of loci is likely to increase the number of incorrect parentage/paternity assignments, the effect of the inclusion of samples with the minimum allowed number genotypes (and their level of polymorphism) should be assessed as well to ascertain that the minimum number of genotypes is sufficient to identify PO pairs and SDO trios with a reasonable power.

In conclusion, we presented a new set of 10 STR loci and one Y chromosome marker to be amplified in two multiplexes in baleen whales. This set of markers serves as an easy, optimized starting point to conduct individual-based and population genetic studies in baleen whales, producing data that later can be shared among research groups. We also provide extended sets of STR loci (from previously published sources) with which to conduct rigorous parentage/paternity assignments along with Python scripts to assess the statistical power of specific sets of STR loci in order to ascertain the power to exclude non-parental genotypes.

## Data availability statement

All scripts are available at https://github.com/MSuarezMenendez/PaternityExclusion. The genotype data are available upon request.

## Author contributions

MSM conducted the data analysis and drafted the manuscript. MB performed DNA extractions, selected markers, optimized PCR reactions, conducted data analysis and drafted the manuscript. PJP designed primers, discussed results and revised all drafts of the manuscripts. LB and ØW provided extracted DNA. TO genotyped the selected markers in DNA samples from different species. MT provided access to genome data. PB, MPHJ, VL, CR, JR, RS, MAS and GV provided samples. All co-authors revised and approved the final manuscript.

## Acknowledgements

MSM was supported by a doctorate fellowship from the University of Groningen. MAS was supported by AZORES 2020, through the EU Fund 01-0145-FEDER-000140. The authors would like to thank Mariel T.I ten Doeschate for samples contribution and W. Hao for her technical assistance.

---

**Table S1.**
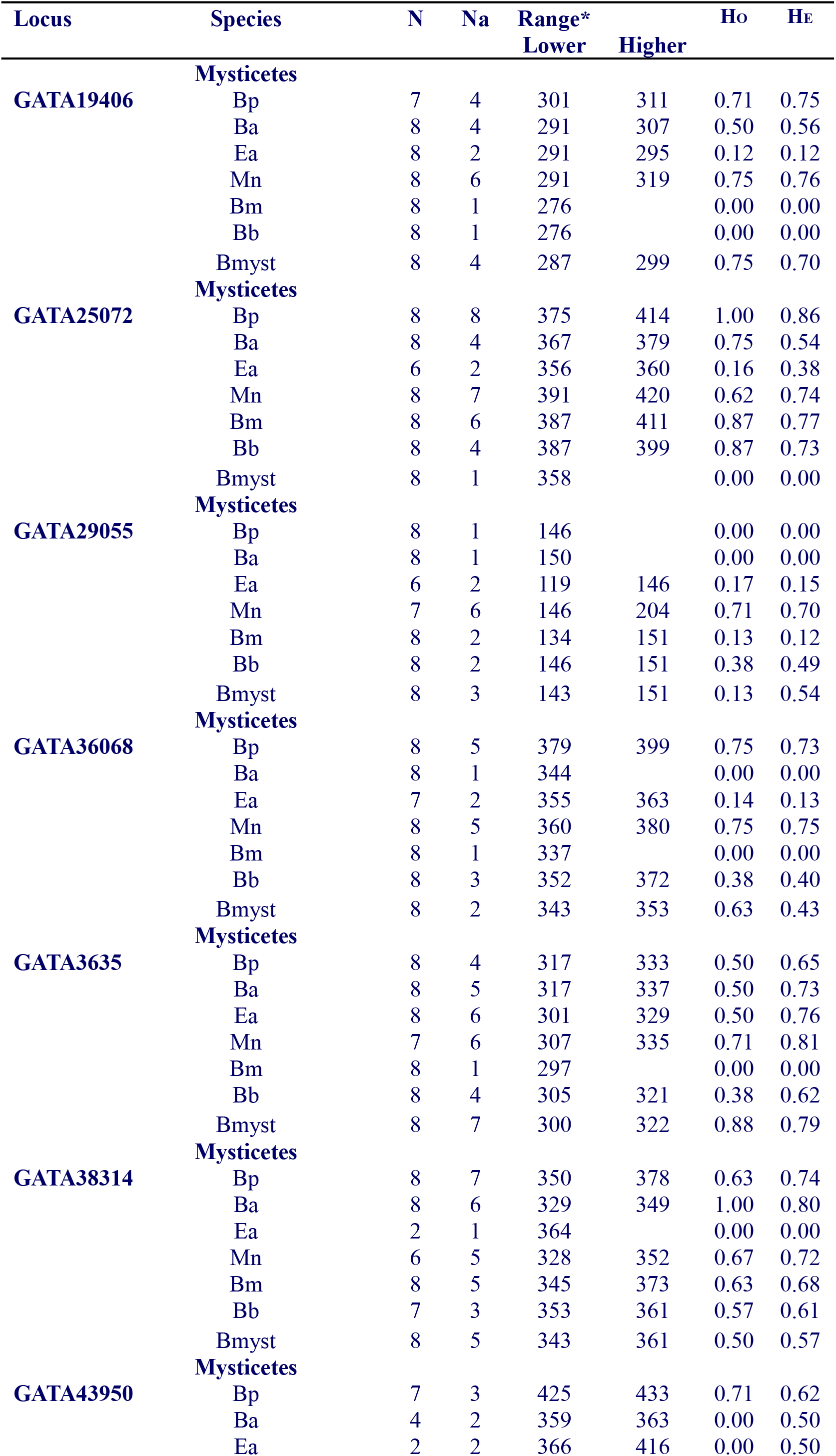
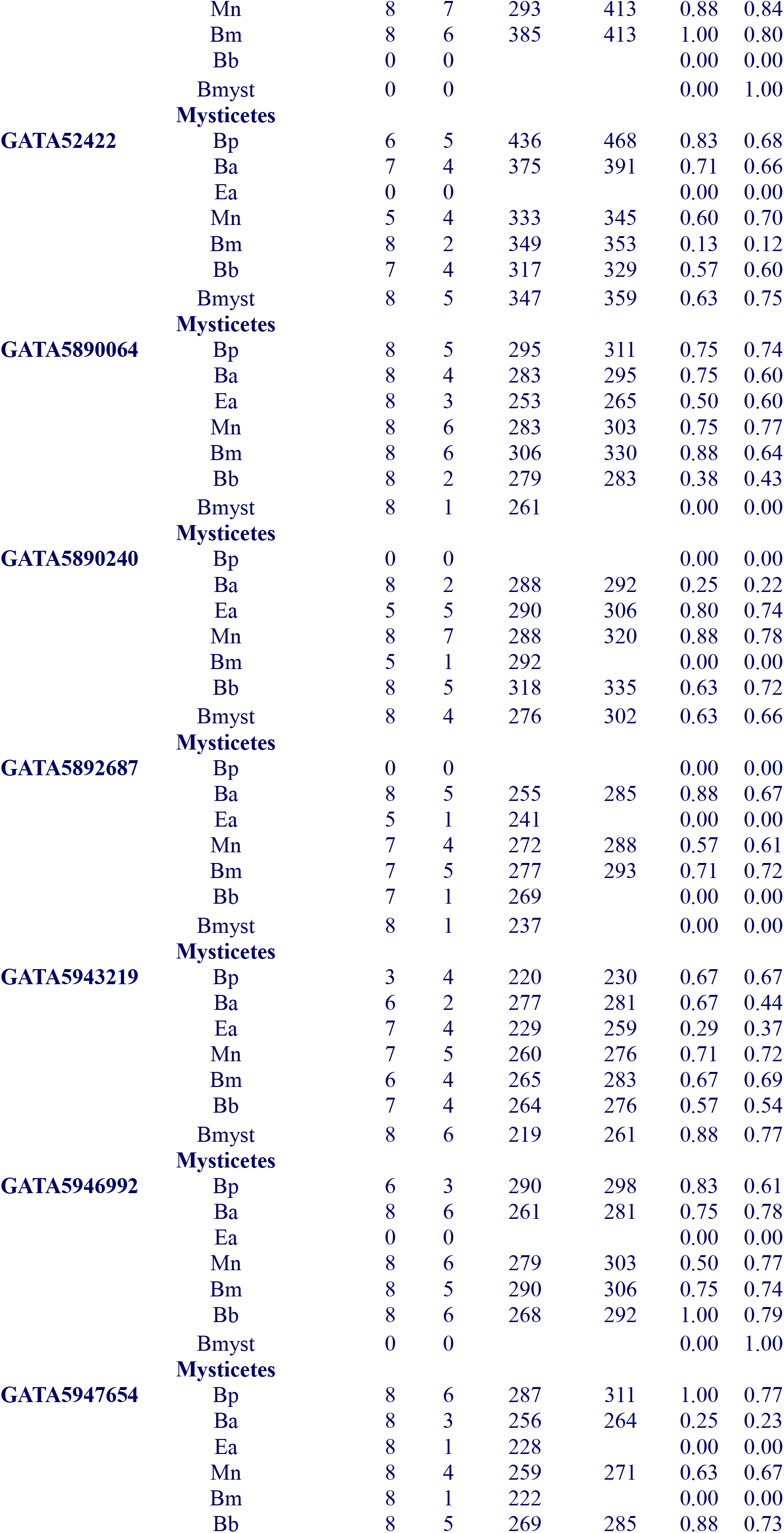
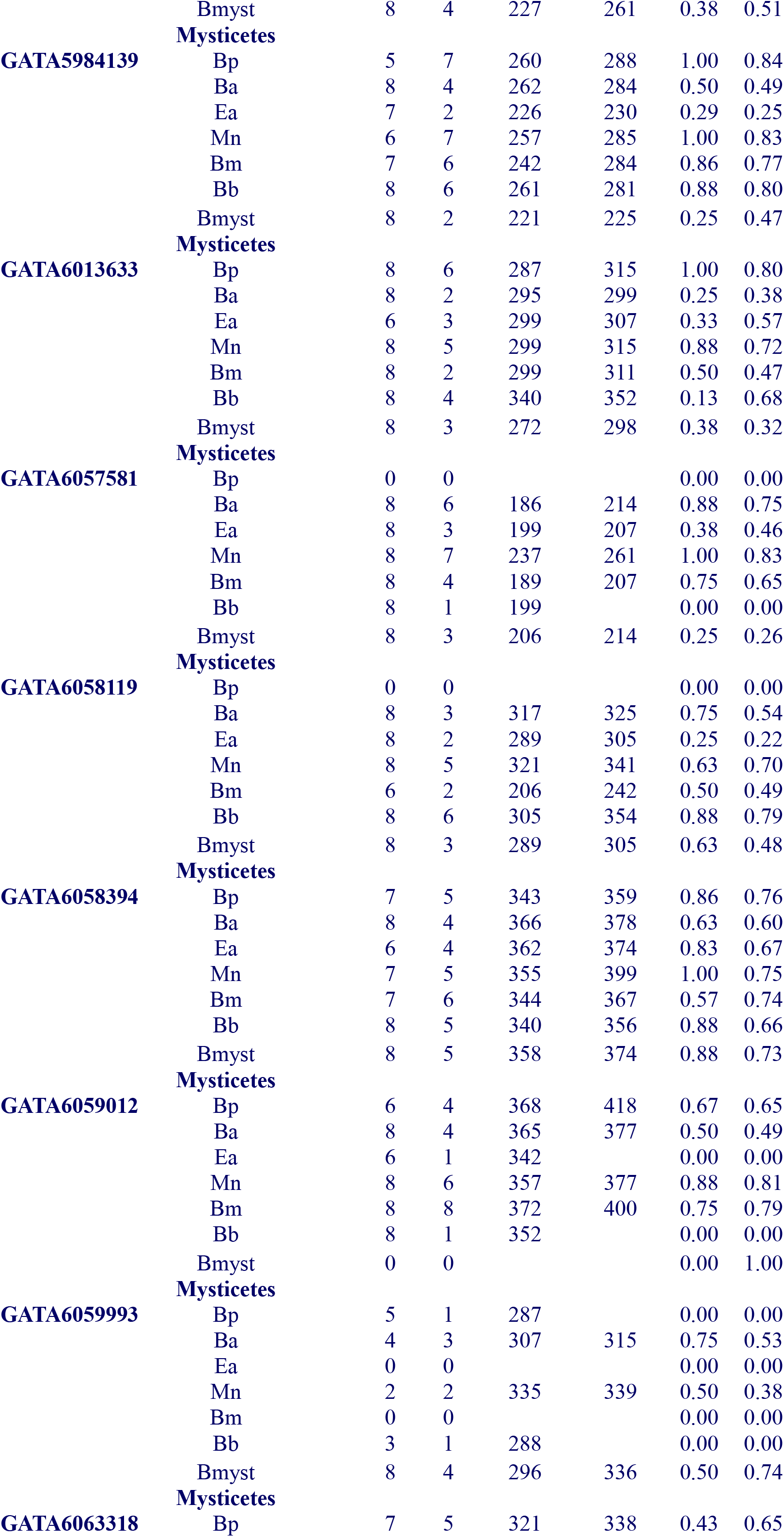
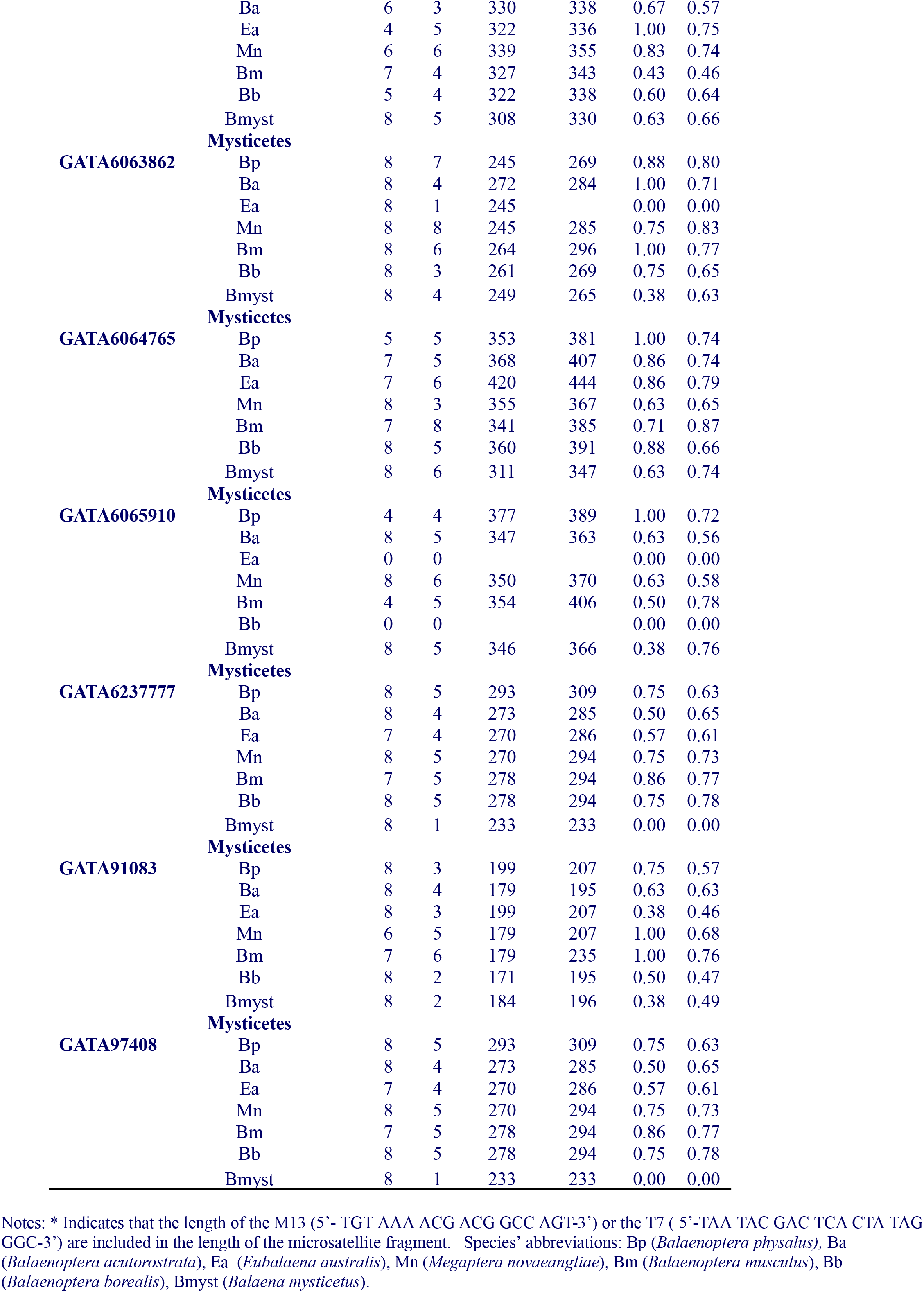
Allele number, size range (bp), observed and expected heterozygosity of 28 novel microsatellite loci in seven Mysticetes species.

**Table S2.** STR loci used for parentage assignment.

| STR loci | Species | $H_E$ | | | | $P_{\text{NON-EXCL}}$ | | | |
| --- | --- | --- | --- | --- | --- | --- | --- | --- | --- |
|  | isolated from | Bp | Mn | Bm | Bmyst | Bp | Mn | Bm | Bmyst |
| AC045<sup>&</sup> | Mn | n/a | 0.74 | n/a | n/a | n/a | 0.52 | n/a | n/a |
| AC082<sup>&</sup> | Mn | n/a | n/a | 0.60 | 0.68 | n/a | n/a | 0.35 | 0.40 |
| AC087<sup>&</sup> | Mn | 0.69 | 0.80 | 0.57 | 0.74 | 0.74 | 0.60 | 0.35 | 0.52 |
| AC137<sup>&</sup> | Mn | n/a | n/a | 0.67 | n/a | n/a | n/a | 0.45 | n/a |
| Bmy16<sup>†</sup> | Bmyst | n/a | n/a | n/a | 0.72 | n/a | n/a | n/a | 0.50 |
| Bmy19<sup>†</sup> | Bmyst | n/a | n/a | n/a | 0.84 | n/a | n/a | n/a | 0.69 |
| Bmy26<sup>†</sup> | Bmyst | n/a | n/a | n/a | 0.91 | n/a | n/a | n/a | 0.80 |
| Bmy33<sup>†</sup> | Bmyst | n/a | n/a | n/a | 0.80 | n/a | n/a | n/a | 0.61 |
| Bmy41<sup>†</sup> | Bmyst | n/a | n/a | n/a | 0.93 | n/a | n/a | n/a | 0.84 |
| Bmy42<sup>†</sup> | Bmyst | n/a | n/a | n/a | 0.80 | n/a | n/a | n/a | 0.63 |
| Bmy58<sup>†</sup> | Bmyst | n/a | n/a | n/a | 0.92 | n/a | n/a | n/a | 0.83 |
| CA128<sup>&</sup> | Bp | n/a | n/a | 0.67 | n/a | n/a | n/a | 0.38 | n/a |
| CA141<sup>&</sup> | Bp | 0.69 | n/a | 0.53 | 0.61 | 0.73 | n/a | 0.32 | 0.33 |
| CA232<sup>&</sup> | Mn | 0.57 | n/a | n/a | 0.76 | 0.83 | n/a | n/a | 0.52 |
| CA234<sup>&</sup> | Mn | 0.81 | 0.73 | 0.71 | n/a | 0.54 | 0.48 | 0.48 | n/a |
| EV001<sup>°</sup> | Pm | 0.84 | 0.63 | n/a | n/a | 0.49 | 0.38 | n/a | n/a |
| EV037<sup>°</sup> | Mn | 0.85 | 0.87 | 0.53 | n/a | 0.46 | 0.74 | 0.33 | n/a |
| EV094<sup>°</sup> | Mn | 0.91 | 0.68 | n/a | n/a | 0.31 | 0.43 | n/a | n/a |
| EV096<sup>°</sup> | Mn | n/a | 0.80 | n/a | n/a | n/a | 0.61 | n/a | n/a |
| GATA028<sup>‡</sup> | Mn | 0.87 | 0.46 | 0.82 | 0.84 | 0.41 | 0.28 | 0.63 | 0.68 |
| GATA053<sup>‡</sup> | Mn | n/a | 0.85 | n/a | n/a | n/a | 0.68 | n/a | n/a |
| GATA098<sup>‡</sup> | Mn | 0.74 | 0.65 | 0.67 | 0.63 | 0.66 | 0.43 | 0.44 | 0.41 |
| GATA19406<sup>α</sup> | Mn | 0.83 | 0.72 | n/a | 0.64 | 0.51 | 0.50 | n/a | 0.35 |
| GATA25072<sup>α</sup> | Mn | 0.87 | 0.84 | n/a | n/a | 0.41 | 0.68 | n/a | n/a |
| GATA38314<sup>α</sup> | Mn | 0.82 | 0.71 | n/a | 0.85 | 0.52 | 0.46 | n/a | 0.69 |
| GATA417<sup>‡</sup> | Mn | 0.86 | 0.86 | 0.87 | n/a | 0.42 | 0.71 | 0.73 | n/a |
| GATA43950<sup>α</sup> | Mn | 0.60 | 0.88 | 0.81 | n/a | 0.79 | 0.75 | 0.63 | n/a |
| GATA52422<sup>α</sup> | Mn | 0.85 | 0.70 | n/a | 0.69 | 0.44 | 0.44 | n/a | 0.44 |
| GATA5947654<sup>α</sup> | Mn | 0.78 | 0.67 | n/a | n/a | 0.59 | 0.41 | n/a | n/a |
| GATA5984139<sup>α</sup> | Mn | 0.83 | 0.84 | n/a | 0.49 | 0.50 | 0.67 | n/a | 0.19 |
| GATA6059012<sup>α</sup> | Mn | 0.75 | n/a | n/a | n/a | 0.65 | n/a | n/a | n/a |
| GATA6063318<sup>α</sup> | Mn | 0.77 | 0.81 | n/a | 0.66 | 0.61 | 0.63 | n/a | 0.42 |
| GATA6063862<sup>α</sup> | Mn | n/a | 0.72 | n/a | n/a | n/a | 0.49 | n/a | n/a |
| GATA6064765<sup>α</sup> | Mn | 0.83 | n/a | n/a | 0.77 | 0.50 | n/a | n/a | 0.58 |
| GATA6237777<sup>α</sup> | Mn | 0.43 | 0.83 | n/a | n/a | 0.91 | 0.65 | n/a | n/a |
| GATA91083<sup>α</sup> | Mn | 0.61 | 0.79 | 0.78 | n/a | 0.80 | 0.59 | 0.58 | n/a |
| GATA97408<sup>α</sup> | Mn | 0.77 | 0.76 | 0.79 | n/a | 0.62 | 0.55 | 0.58 | n/a |
| GT011<sup>□</sup> | Mn | 0.83 | 0.80 | n/a | 0.60 | 0.50 | 0.60 | n/a | 0.33 |
| GT015<sup>α</sup> | Mn | n/a | 0.78 | n/a | n/a | n/a | 0.57 | n/a | n/a |
| GT023<sup>#</sup> | Mn | 0.78 | 0.80 | 0.83 | 0.74 | 0.60 | 0.60 | 0.64 | 0.51 |
| GT101<sup>#</sup> | Mn | n/a | 0.71 | 0.69 | n/a | n/a | 0.49 | 0.46 | n/a |
| GT129<sup>&</sup> | Bp | n/a | n/a | n/a | 0.62 | n/a | n/a | n/a | 0.34 |
| GT195<sup>#</sup> | Mn | n/a | 0.62 | n/a | n/a | n/a | 0.32 | n/a | n/a |
| GT211<sup>#</sup> | Mn | 0.80 | 0.79 | n/a | n/a | 0.56 | 0.58 | n/a | n/a |
| GT271<sup>#</sup> | Mn | 0.64 | 0.51 | 0.53 | n/a | 0.75 | 0.31 | 0.28 | n/a |
| GT310<sup>#</sup> | Mn | 0.71 | n/a | 0.51 | n/a | 0.67 | n/a | 0.23 | n/a |
| GT541<sup>&</sup> | Mn | n/a | n/a | 0.73 | n/a | n/a | n/a | 0.50 | n/a |
| GT575<sup>#</sup> | Mn | 0.69 | 0.73 | 0.63 | n/a | 0.72 | 0.52 | 0.35 | n/a |
| TAA031<sup>‡</sup> | Mn | n/a | 0.78 | n/a | n/a | n/a | 0.57 | n/a | n/a |
| TAA023<sup>‡</sup> | Mn | 0.76 | n/a | n/a | n/a | 0.62 | n/a | n/a | n/a |
| TR3G2* | Eg | n/a | n/a | 0.76 | n/a | n/a | n/a | 0.55 | n/a |

Notes: Mn denotes *Megaptera novaeangliae*, Bmyst denotes *Balaena mysticetus* and Bm, *Balaenoptera musculus*, Bp, *Balaenoptera physalus*, Eg, *Eubalaena glacialis*, Pm, *Physeter macrocephalus*. H<sub>E</sub> denotes expected heterozygosity and P<sub>NON-EXCL</sub>, the non-exclusion probability of the first parent. References; <sup>□</sup>(Bérubé et al. 1998); <sup>#</sup>(Bérubé et al. 2000); <sup>&</sup>(Bérubé et al. 2005); <sup>\*</sup>(Frasier et al. 2006) ; <sup>¶</sup>(Huebinger et al. 2008); <sup>‡</sup>(Palsbøll et al. 1997); <sup>“</sup>This study; <sup>°</sup>(Valsecchi and Amos 1996)

**Table S3.**
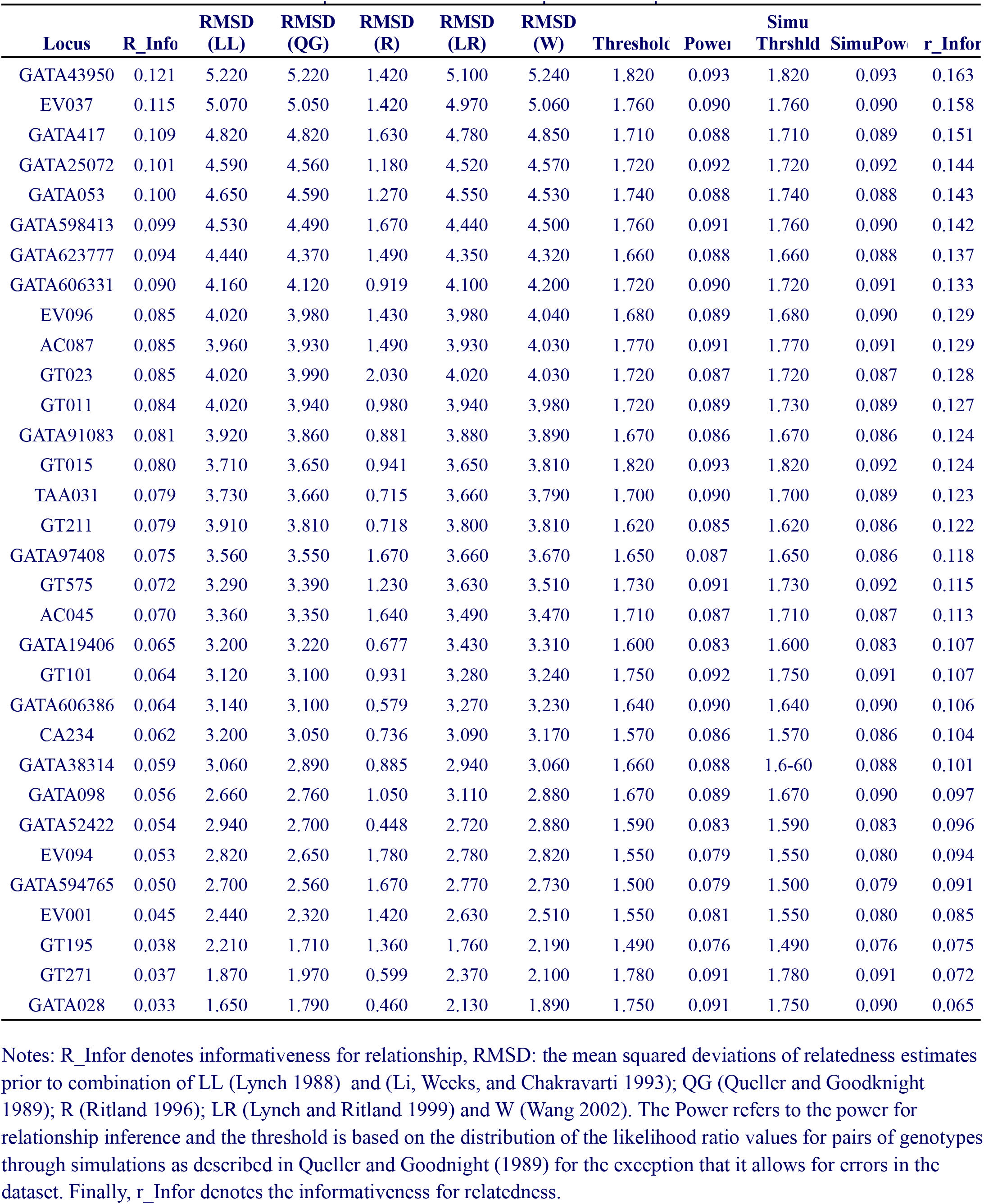
Informativeness estimates and power to detect relationships in the humpback whale dataset

| Locus | R_Info | RMSD (LL) | RMSD (QG) | RMSD (R) | RMSD (LR) | RMSD (W) | Threshold | Power | Simu Thrshld | SimuPow | r_Infor |
| --- | --- | --- | --- | --- | --- | --- | --- | --- | --- | --- | --- |
| GATA43950 | 0.121 | 5.220 | 5.220 | 1.420 | 5.100 | 5.240 | 1.820 | 0.093 | 1.820 | 0.093 | 0.163 |
| EV037 | 0.115 | 5.070 | 5.050 | 1.420 | 4.970 | 5.060 | 1.760 | 0.090 | 1.760 | 0.090 | 0.158 |
| GATA417 | 0.109 | 4.820 | 4.820 | 1.630 | 4.780 | 4.850 | 1.710 | 0.088 | 1.710 | 0.089 | 0.151 |
| GATA25072 | 0.101 | 4.590 | 4.560 | 1.180 | 4.520 | 4.570 | 1.720 | 0.092 | 1.720 | 0.092 | 0.144 |
| GATA053 | 0.100 | 4.650 | 4.590 | 1.270 | 4.550 | 4.530 | 1.740 | 0.088 | 1.740 | 0.088 | 0.143 |
| GATA598413 | 0.099 | 4.530 | 4.490 | 1.670 | 4.440 | 4.500 | 1.760 | 0.091 | 1.760 | 0.090 | 0.142 |
| GATA623777 | 0.094 | 4.440 | 4.370 | 1.490 | 4.350 | 4.320 | 1.660 | 0.088 | 1.660 | 0.088 | 0.137 |
| GATA606331 | 0.090 | 4.160 | 4.120 | 0.919 | 4.100 | 4.200 | 1.720 | 0.090 | 1.720 | 0.091 | 0.133 |
| EV096 | 0.085 | 4.020 | 3.980 | 1.430 | 3.980 | 4.040 | 1.680 | 0.089 | 1.680 | 0.090 | 0.129 |
| AC087 | 0.085 | 3.960 | 3.930 | 1.490 | 3.930 | 4.030 | 1.770 | 0.091 | 1.770 | 0.091 | 0.129 |
| GT023 | 0.085 | 4.020 | 3.990 | 2.030 | 4.020 | 4.030 | 1.720 | 0.087 | 1.720 | 0.087 | 0.128 |
| GT011 | 0.084 | 4.020 | 3.940 | 0.980 | 3.940 | 3.980 | 1.720 | 0.089 | 1.730 | 0.089 | 0.127 |
| GATA91083 | 0.081 | 3.920 | 3.860 | 0.881 | 3.880 | 3.890 | 1.670 | 0.086 | 1.670 | 0.086 | 0.124 |
| GT015 | 0.080 | 3.710 | 3.650 | 0.941 | 3.650 | 3.810 | 1.820 | 0.093 | 1.820 | 0.092 | 0.124 |
| TAA031 | 0.079 | 3.730 | 3.660 | 0.715 | 3.660 | 3.790 | 1.700 | 0.090 | 1.700 | 0.089 | 0.123 |
| GT211 | 0.079 | 3.910 | 3.810 | 0.718 | 3.800 | 3.810 | 1.620 | 0.085 | 1.620 | 0.086 | 0.122 |
| GATA97408 | 0.075 | 3.560 | 3.550 | 1.670 | 3.660 | 3.670 | 1.650 | 0.087 | 1.650 | 0.086 | 0.118 |
| GT575 | 0.072 | 3.290 | 3.390 | 1.230 | 3.630 | 3.510 | 1.730 | 0.091 | 1.730 | 0.092 | 0.115 |
| AC045 | 0.070 | 3.360 | 3.350 | 1.640 | 3.490 | 3.470 | 1.710 | 0.087 | 1.710 | 0.087 | 0.113 |
| GATA19406 | 0.065 | 3.200 | 3.220 | 0.677 | 3.430 | 3.310 | 1.600 | 0.083 | 1.600 | 0.083 | 0.107 |
| GT101 | 0.064 | 3.120 | 3.100 | 0.931 | 3.280 | 3.240 | 1.750 | 0.092 | 1.750 | 0.091 | 0.107 |
| GATA606386 | 0.064 | 3.140 | 3.100 | 0.579 | 3.270 | 3.230 | 1.640 | 0.090 | 1.640 | 0.090 | 0.106 |
| CA234 | 0.062 | 3.200 | 3.050 | 0.736 | 3.090 | 3.170 | 1.570 | 0.086 | 1.570 | 0.086 | 0.104 |
| GATA38314 | 0.059 | 3.060 | 2.890 | 0.885 | 2.940 | 3.060 | 1.660 | 0.088 | 1.6-60 | 0.088 | 0.101 |
| GATA098 | 0.056 | 2.660 | 2.760 | 1.050 | 3.110 | 2.880 | 1.670 | 0.089 | 1.670 | 0.090 | 0.097 |
| GATA52422 | 0.054 | 2.940 | 2.700 | 0.448 | 2.720 | 2.880 | 1.590 | 0.083 | 1.590 | 0.083 | 0.096 |
| EV094 | 0.053 | 2.820 | 2.650 | 1.780 | 2.780 | 2.820 | 1.550 | 0.079 | 1.550 | 0.080 | 0.094 |
| GATA594765 | 0.050 | 2.700 | 2.560 | 1.670 | 2.770 | 2.730 | 1.500 | 0.079 | 1.500 | 0.079 | 0.091 |
| EV001 | 0.045 | 2.440 | 2.320 | 1.420 | 2.630 | 2.510 | 1.550 | 0.081 | 1.550 | 0.080 | 0.085 |
| GT195 | 0.038 | 2.210 | 1.710 | 1.360 | 1.760 | 2.190 | 1.490 | 0.076 | 1.490 | 0.076 | 0.075 |
| GT271 | 0.037 | 1.870 | 1.970 | 0.599 | 2.370 | 2.100 | 1.780 | 0.091 | 1.780 | 0.091 | 0.072 |
| GATA028 | 0.033 | 1.650 | 1.790 | 0.460 | 2.130 | 1.890 | 1.750 | 0.091 | 1.750 | 0.090 | 0.065 |

Notes: R\_Info denotes informativeness for relationship, RMSD: the mean squared deviations of relatedness estimates prior to combination of LL (Lynch 1988) and (Li, Weeks, and Chakravarti 1993); QG (Queller and Goodnight 1989); R (Ritland 1996); LR (Lynch and Ritland 1999) and W (Wang 2002). The Power refers to the power for relationship inference and the threshold is based on the distribution of the likelihood ratio values for pairs of genotypes through simulations as described in Queller and Goodnight (1989) for the exception that it allows for errors in the dataset. Finally, r\_Info denotes the informativeness for relatedness.

